# Phenotypic and genomic evidence for transparent cellulose, metabolic diversity, and stable cellulose production in the Acetobacteraceae

**DOI:** 10.1101/2023.08.21.554206

**Authors:** Kevin W. Keating, Elizabeth M. van Zyl, Joseph H. Collins, Carter Nakagawa, Sarah J. Weintraub, Jeannine M. Coburn, Eric M. Young

**Author notes:** co-first authors (Kevin W. Keating, Elizabeth M. van Zyl).

## Abstract

The Acetobacteraceae are a family of microbes that use sugars from fruits, beverages and fermented foods to overproduce bacterial nanocellulose (BNC), a living material with broad applications in medicine and industry. Yet, the family has few complete, contiguous genome sequences available. Here, three different strains – a high production strain NQ5, a metabolic engineering host NCIB 8034, and a new isolate DS12 from kombucha were characterized and complete *de novo* genomes assembled. Initial growth and yield experiments reveal a diversity of carbon source utilization profiles and BNC production rates, with NQ5 achieving the highest yield on glucose and DS12 having the narrowest utilization profile. All strains synthesize optically clear BNC. Genomic evidence assigns the DS12 isolate to *Komagataeibacter nataicola,* reassigns NCIB 8034 from *Komagataeibacter xylinus* to *Komagataeibacter oboediens,* and supports NQ5 as *Novacetimonas hansenii.* The *bcs* gene clusters that encode BNC synthesis are also diverse. The highest producing strain, *N. hansenii* NQ5, has fewer *bnc* copies than *K. oboediens* NQ5, indicating that copy number does not explain high productivity. Analysis also reveals the type and frequency of mobile genetic elements. Notably, *N. hansenii* NQ5 has a paucity of transposons relative to other strains, which could explain the BNC production stability of *N. hansenii* NQ5 in culture. Thus, this work argues that Acetobacteraceae are metabolically diverse, and provides genomic evidence explaining beneficial BNC production characteristics of *N. hansenii* NQ5. Therefore, this work provides evidence for selection of appropriate BNC production strains.

**IMPORTANCE:** Bacterial cellulose is an important material for biomedical applications like wound dressings. This study defines important characteristics of microbes that produce bacterial cellulose, namely their ability to process different sugars and features of their genomes that make cellulose yield more consistent. These findings will aid in the development of better bacterial cellulose production processes.

## INTRODUCTION

Acetobacteraceae is a family of Alphaproteobacteria that overproduce bacterial nanocellulose (BNC), which has applications in biomedical^1^, electronics^2^, and food^3^ industries. These bacteria grow on inexpensive, complex carbon sources derived from fruit and sugarcane, such as molasses^4,5^, and are notable members of microbial communities in cultured products like kombucha, nata de coco, and vinegar. Many of these strains are now sequenced, which enables characterization of the genome architecture and copy number of the BNC synthesis operons across the family^6–8^. These analyses are key to studying the evolutionary history of BNC synthesis and understanding how strain genetics inform BNC yield and properties for bioprocessing. Even so, many of the available genomes are fragmentary and incomplete, particularly for some of the most commercially relevant strains. Recently, we developed a workflow to accurately sequence and assemble yeast genomes using short and long reads. Here, we apply a similar strategy to three diverse strains of Acetobacteraceae and correlate the resultant genomes to observed growth and BNC production phenotypes.

Over the past decade, the number of described Acetobacteraceae strains has rapidly increased which has driven phylogenetic reclassification of the family. Most BNC overproducing strains were grouped into the *Komagataeibacter* genus, but recent whole genome analysis has argued for some to be placed in a new genus, *Novacetimonas*^9^. This includes the most robust BNC producing strain, *Novacetimonas hansenii* NQ5^10^. *Novacetimonas* also includes the recent validly described species *N. cocois*^11^, *N. maltaceti*^12^, and *N. pomaceti*^13^, while *Komagataeibacter* retains *K. xylinus* (formerly *Gluconacetobacter xylinus*) as well as recent validly described species such as *K. oboediens*, *K. melaceti*^14^, *K. melomensus*^14^, *K. kakiaceti*^15^, *K. medellinensis*^16^, and *K. diospyri*^17^. This demonstrates the need for high-quality, strain-specific genomes to accurately classify strains and contextualize observed phenotypes within the phylogeny.

BNC is an attractive material for a variety of reasons. It has high purity – unlike cellulose derived from plants, it does not have hemicellulose, lignin, and pectin contaminants^18^. It is biocompatible and generally regarded as safe (GRAS). It is also flexible, highly crystalline, strong, and porous which combine to give it high structural integrity and good mass transfer properties. Therefore, BNC derived materials are used in wound healing and drug delivery^19^. Recently, an approach to make optically clear BNC with strain *N. hansenii* NQ5 was reported, opening up new possibilities for BNC use in biomedical applications^20^.

BNC is synthesized by *bcsA*, an inner membrane protein that uses cyclic di-GMP as a cofactor to assemble UDP-glucose units into a β(1,4)-glucan fibril. Brandão et al. describe four types of gene clusters present in *Gluconacetobacter*, *Komagataeibacter*, and *Novacetimonas* that contain additional genes beyond *bcsA*. The gene *bcsB* combines with *bcsA* to form the catalytically active bcs core, and is essential for cellulose synthesis *in vitro*^21^. Both *bcsA* and *bcsB* are present in all gene clusters, and in some cases are expressed as a single coding sequence. The nascent cellulose fibril is assembled into fibers by outer membrane proteins *bcsD* and *bcsC,* which are present in nearly all *Komagataeibacter* and *Novacetimonas* species as a *bcsABCD* cluster, often preceded by accessory genes *bcsZ* and *bcsH.* It is thought that the copy number and efficiency of these enzymes influence BNC crystallinity and yield. This is also affected by the age of the culture, as often in serial liquid subculture BNC synthesis is lost, hinting at a common regulatory process or genome instability that controls BNC production. Three additional gene clusters, *bcsABXYC*, *bcsABC*, and *bcsAB*, are widespread among these cellulose-producing genera. Though the roles and regulation of each gene cluster have yet to be determined, it has been proposed that the *bcsABXYC* cluster produces an acylated form of BNC^22^.

The commercial relevance of this family and increasing genomic understanding has motivated metabolic modeling and genetic engineering. Yadav *et al.* engineered *K. xylinus* NCIB 8034 (formerly *G. xylinus*), showing that *bcsA* could accept UDP-glcNAc and synthesize a cellulose-chitin hybrid material^23^. Florea *et al.* domesticated a strain of *K. rhaeticus* and demonstrated the function of genetic circuits in the strain.^24^ We have shown that many genetic parts from *E. coli* can be functional in the novel strain isolated in this report, DS12.^25^ Finally, a recent report described the first genome-scale metabolic model for Acetobacteraceae, focusing on *K. xylinus* DSM 2325.^26^ Thus, it is now possible to genetically engineer Acetobacteraceae for the control and modification of BNC, yet this cannot be done precisely without quality genome sequences, which are not yet available.

In this work, we studied three strains of significant interest to BNC production. We obtained phenotypic data and high-quality genome sequences for each strain. The results support that *N. hansenii* NQ5 should be the ideal chassis for future BNC production efforts because of its genome stability, high BNC yield on glucose, ability to synthesize optically clear BNC, and broad carbon source palette.

## RESULTS

### Strain selection and isolation

We selected the emerging production host NQ5 (ATCC #53582) and the previously engineered NCIB 8034 (NRC 6018, ATCC #10245) because they represent different branches of Acetobacteraceae and were isolated from different sources. NQ5 was isolated from a sugar mill in North Queensland, Australia. Though two genomes for NQ5 are currently available on the NCBI genome database (accessions GCF_001645815.1 and GCF_003416815.1), both assemblies are highly fragmented draft genomes due to short read-only sequencing. NCIB 8034 is of unknown provenance and there is no genome available. To obtain a representative strain from cultured products, strain DS12 was isolated from kombucha (**Figure 1 A-E**, **Methods**). In brief, cellulose producing colonies sourced from a home-brewed kombucha pellicle were selected from agar plates for subculture in Hestrin-Schramm media (HS) and screened for retention of cellulose production after serial subculture. The resulting isolated strain was named DS12. The NQ5 strain has been used in the laboratory and available in culture collection since at least 1987^27^, and the NCIB 8034 strain was available in a culture collection as cited in a patent in 1966^28^, compared to the isolation of DS12 in 2018. Together, these three strains represent different lineages and sources, providing diverse backgrounds to investigate the similarities and differences in the Acetobacteraceae.

**Figure 1:**
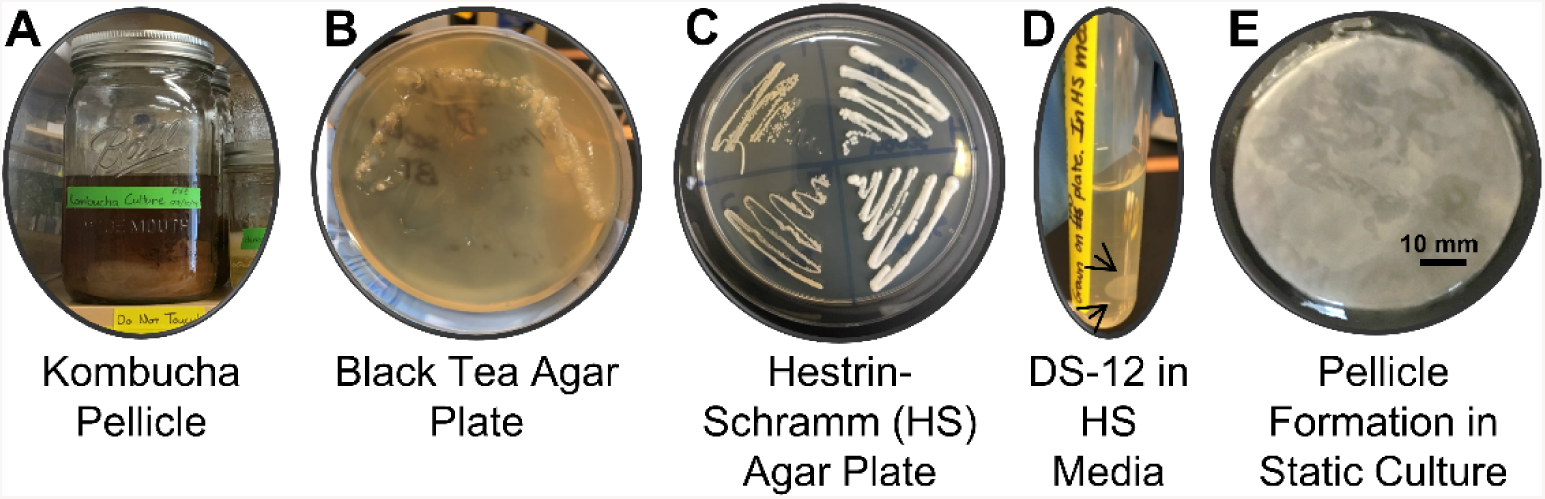
Isolation of DS-12. **A**. The kombucha pellicle from which DS-12 was isolated. **B**. Swab from pellicle was streaked onto black tea agar plate. **C**. Growth was then swabbed onto HS agar plates. **D.** Colonies were inoculated into HS media. **E.** Cellulose formation of DS-12 was confirmed in static culture.

### Phenotypic characterization – Growth

First, we phenotypically characterized the three strains for their carbon source utilization profile, BNC yield, and BNC transparency. To obtain a carbon source profile, we modified the defined Hestrin and Schramm^29^ media by substituting 10 carbon sources besides glucose – sucrose, fructose, galactose, mannose, mannitol, N-acetyl-D-glucosamine (GlcNAc), glucosamine (GlcN), xylose, arabinose, and arabitol. The hexoses, pentoses, disaccharides, and sugar alcohols are common constituents of biomass and many have been used previously as carbon sources for cellulose production.^30–36^ GlcNAc and glucosamine represent sugar monomers with chemically functional side groups for incorporation and subsequent chemical modification.^23,37,38^ Then, we grew each strain on this modified HS medium and measured the optical density (OD600) over time. Maximum growth rate (µmax) values (**Figure 2A, Supplementary Table S1**) were calculated using the maximum slope from the optical density measurements. Growth was observed on every carbon source except GlcN, consistent with its known toxicity^39^. Both NQ5 and NCIB 8034 exhibited maximum growth rates of 0.005 or more on all carbon sources, save NQ5 on mannose, which none of the strains grew well on. However, DS12 only achieved growth faster than 0.005 on fructose, glucose, and mannitol. The fast growth on glucose and fructose, and limited growth on sucrose, may be an adaptation to kombucha culture. It has been shown in the kombucha symbiotic culture of yeast and bacteria (SCOBY) that yeasts with extracellular invertase activity catabolize sucrose into fructose and glucose^40^. Thus, microbes would have access to abundant glucose and fructose and have no pressure to retain sucrose metabolism. This also may explain the relatively poor growth on all other carbon sources for DS12 in general. The fast growth of all strains on mannitol versus mannose is interesting to note since they are catabolized through different pathways. A complete PEP system in *Komagataeibacter* has not been clearly established, but mannitol has been reported to be transported into *K. nataicola* RZS01 via the phosphotransferase system, being phosphorylated into mannitol-1-phosphate in the process.^41^ It is then isomerized into fructose-6-phosphate and likely is catabolized similarly to fructose, perhaps explaining the similar growth rates.

**Figure 2:**
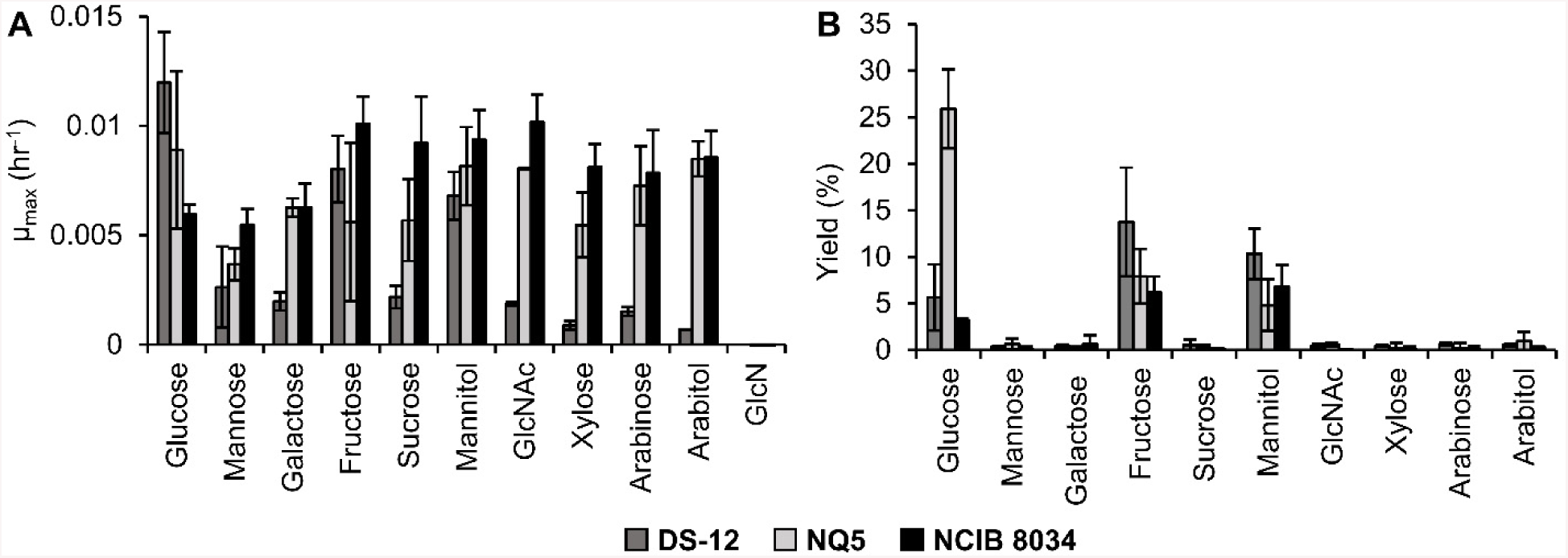
Maximum growth rate (**A**) and cellulose yield (**B**) of DS12, NCIB 8034, and NQ5 grown on various carbon sources.

### Phenotypic characterization – BNC yield, transparency, and absorption

We then measured BNC production on all carbon sources. There was observable BNC production for most carbon sources. Using **Equation 1** (**Methods**), cellulose yield was determined relative to carbon source mass (**Figure 2B, Supplementary Table S2**). Consistent with literature, DS12, NQ5, and NCIB 8034 were found to support high cellulose yield formation when grown with fructose, mannitol, and glucose as a carbon source^42^. NQ5 produced the highest cellulose yield of 25.9% when glucose was used as a carbon source. NCIB 8034 known to prefer mannitol over glucose as the primary carbon source^7,43^, which is consistent with our findings where 3.2% BC yield was observed using glucose, while 6.8% BC yield was observed using mannitol. NCIB 8034 also consistently exhibited higher cellulose yield production when using fructose as a carbon source (6.2%). The carbon sources that resulted in highest BC yield by DS12 were fructose and mannitol, with a yield of 13.8% and 10.3%, respectively. Yet, significant BNC yields were only observed for fructose, glucose, and mannitol. BNC biosynthesis depends on UDP-glucose, and in turn glucose. Therefore, the results appear to show that gluconeogenesis is not robust in these bacteria, as they are able to grow but not produce much BNC on carbon sources that do not have a metabolic path through glucose. Of the five carbon feeds, the highest yield was on arabitol, although the productivity was much less.

The transparency of the BNC products on all sugars was evaluated by overall light transmission in the visible light region using a UV-Vis spectrophotometer (**Methods**). All strains could synthesize transparent BNC, but arabitol particularly resulted in transparent BNC (**Figure 3**). The impact of arabitol and glucose cofeeding on BNC material properties for NQ5 is discussed in more detail in our recent report^44^. These results demonstrate that optically clear properties can be achieved across the Acetobacteraceae, indicating that culture conditions and perhaps synthesis rate are the key factors in transparent cellulose synthesis^45^.

**Figure 3:**
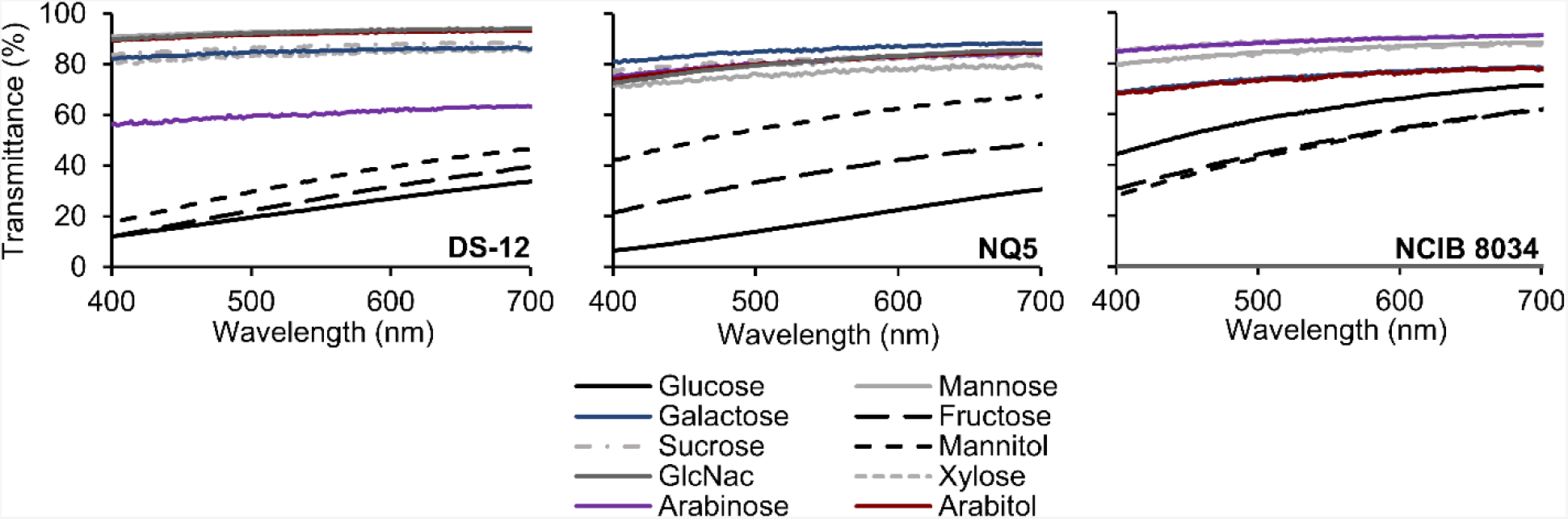
Transparency of the bacterial cellulose produced by D2-12, NQ5, and NCIB 8034 when grown on various carbon sources.

**Figure 4:**
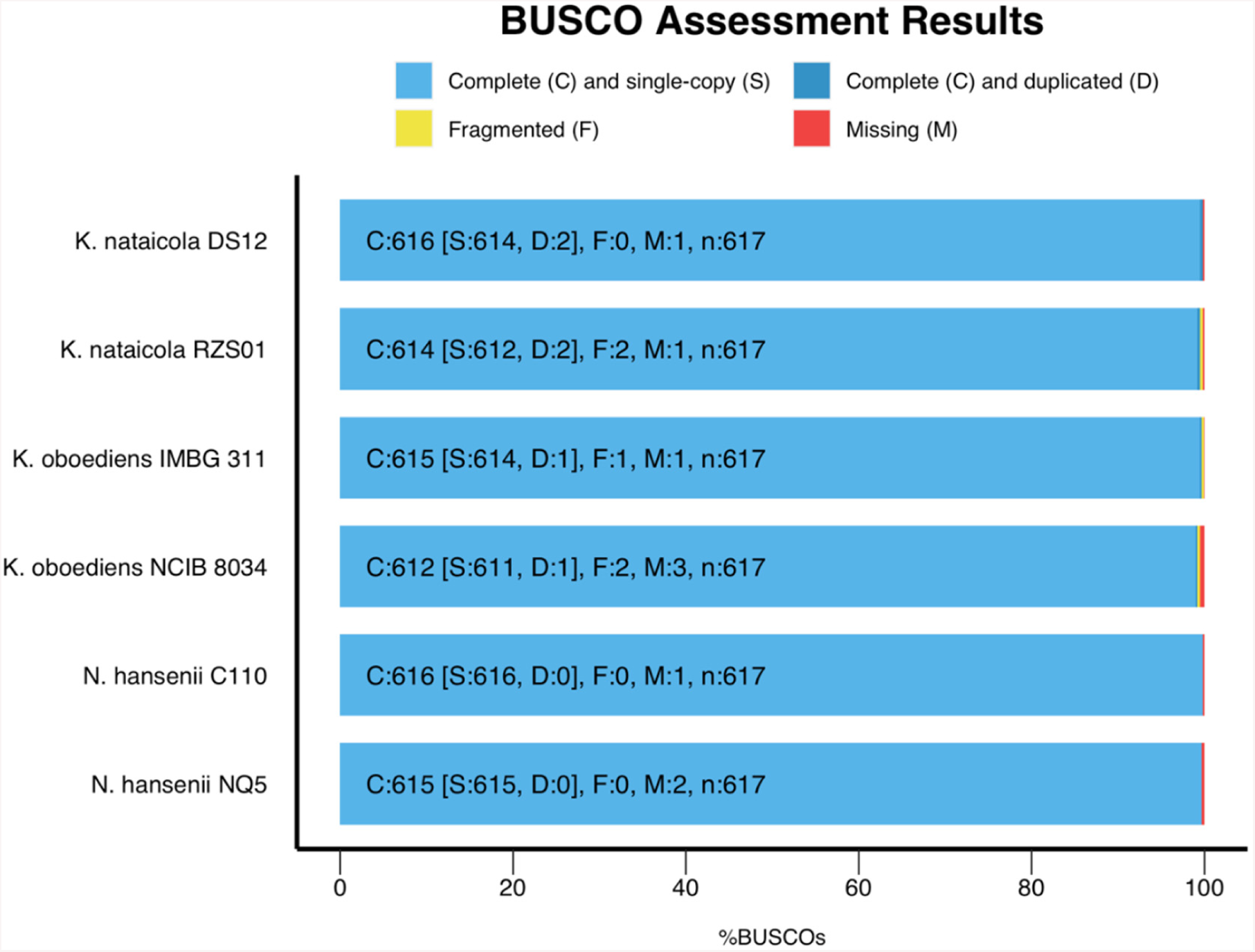
The novel genomes for *K. nataicola* DS12, *K. oboediens* NCIB 8034, *N. hansenii* NQ5 and NCBI representative genomes for each species were assessed for completeness using BUSCO. All genomes were over 99% complete, indicating that the novel genomes are of comparable completeness to the reference genomes.

**Figure 5:**
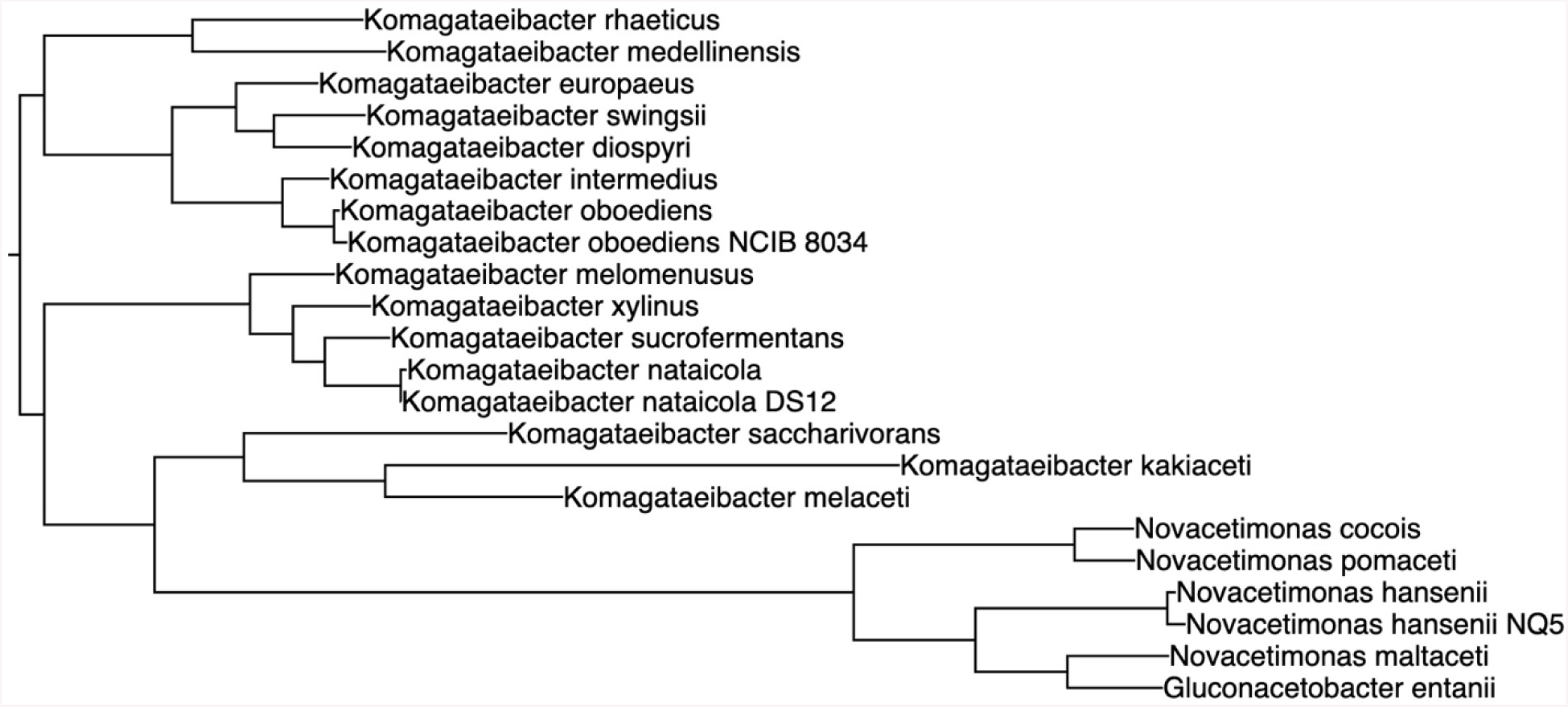
A phlyogenetic tree of the Komagataeibacter and Novacetimonas genera was constructed using Orthofinder. The novel isolate DS12 was assigned to K.nataicola, and NQ5 was confirmed to belong to N.hansenii. The NCIB 8034 strain, deposited to ATCC as Gluconacetobacter xylinus, was determined to be most closely related to the K.oboediens reference strain.

### Whole genome sequencing, assembly, and completeness

Illumina and Oxford Nanopore Next Generation Sequencing (NGS) technologies were employed to sequence NQ5, NCIB 8034, and DS12 (**Methods**). Estimated sequencing depth was determined by multiplying the number of reads by the average read length and dividing by the assembly size, using a custom script. Sequencing depth was greater than 200× for all strains. The bacterial genome assembler Unicycler was then used to assemble the reads into a whole genome sequence. Summary metrics for the sequencing runs and assemblies are included in Error! Reference source not found.

The initial assemblies were uploaded to NCBI and the closest representative genome was identified. The closest available genome to DS12 was *K. nataicola* RZS01, the closest genome to NCIB 8034 was *K. oboediens* IMBG 311, and the closest genome to NQ5 was *N. hansenii* C110. These were used as comparisons throughout the study and as a benchmark for quality.

Genome quality was assessed using QUAST^46^. A complete genome assembly contains a number of contigs equal to the number of replicons. For the bacteria studied here, the number of replicons is expected to be one (chromosome) plus the number of plasmids. Though we do not have molecular evidence for the size and number of plasmids in each strain, replicative element (dnaA and repA) homologs were identified on each contig assigned to a replicon. Assembly metrics for the genomes generated in this study, as well as the NCBI representative genome for each species, are reported in Error! Reference source not found.. The genome assemblies generated here were therefore deemed to be complete.

Compared to complete representative genomes for *K. nataicola* and *N. hansenii*, the number of replicons are reasonable. The number of replicons for *K. oboediens* NCIB 8034 was comparable to the *K. nataicola* strains, but much lower than the reference assembly. This is because the *K. oboediens* IMBG 311 genome is a highly fragmented “scaffold” level assembly. The assembly for NCIB 8034 generated here is therefore more complete than the current representative genome for the *K. oboediens* species.

The percent G+C of each new assembly was expected to be within 1% of the species representative genome^47^. The percentage point differences reported here were *K. nataicola* +0.42, *K. oboediens* –0.37, and *N. hansenii* –0.11, all within the expected range. The largest contig (chromosome) length for *K. nataicola* and *N. hansenii* were close to those of the representative assemblies. The length for the *K. oboediens* representative longest contig was implausibly short, so the two lengths were not comparable.

The BUSCO^48^ quality assessment tool was also used to measure completeness. BUSCO evaluates single-copy orthologs which are nearly universal within a particular taxonomic clade and reports their presence or absence. The genome assemblies generated in this study, as well as representative genomes for each species, were assessed for BUSCO completeness using the Rhodospiralles clade reference genes (617 genes included in the set). Completeness scores are shown in **Figure**. All genomes generated in this study were over 99% complete and appeared comparable to the reference assemblies.

### Phylogenetic placement of strains

The species for the NQ5 and NCIB 8034 strains are reported by ATCC as *Gluconacetobacter hansenii* and *Gluconacetobacter xylinus* (accessed January 17, 2023). Several changes have occurred within the *Gluconacetobacter* genus recently, including the creation of the *Komagataeibacter* genus^49,50^ and the creation of the *Novacetimonas* genus^9^. It is therefore warranted to reassess the taxonomic identification of these strains. Additionally, the DS12 strain is a novel isolate, therefore a phylogenetic placement is appropriate in this study. An initial phylogenetic tree was constructed with OrthoFinder which included all species from the genera *Gluconacetobacter*, *Komagataeibacter*, and *Novacetimonas* (Error! Reference source not found.). A notable result of the inclusion of *Gluconacetobacter* is that the *Gluconacetobacter entanii* AV429 strain clearly belongs in the *Novacetimonas* genus. As discussed by Jelenko et al., this is likely to occur if the AV429 strain is accepted as the neotype strain of *G. entanii*, whose type strain was lost as an unrecoverable stock^51^. For the purposes of this study, *G. entanii* was included within the *Novacetimonas* clade without renaming. All three strains of interest, however, clearly fall within *Komagataeibacter* and *Novacetimonas*, so a phylogenetic tree containing only these genera was generated to better show the placement of the DS12, NQ5, and NCIB 8034 strains (**Figure, Supplementary Table S5**). The minimum percentage of genes assigned to orthogroups was 90.4% (*Komagataeibacter kakiaceti*), with a median of 97.3%.

The novel DS12 strain is placed most closely to *Komagataeibacter nataicola*. This phylogenetic analysis based upon all protein sequences in the genome is confirmed in the nucleotide space by average nucleotide identity (ANI) analysis^52^, which calculates 99.7% identity to the RZS01 strain of *K. nataicola* (Error! Reference source not found.). The next closest species to DS12 by ANI is *K. sucrofermentans* LMG 18788 at 92.6%. As the species identity threshold by ANI is typically set at 96%^53^, the DS12 strain can be unambiguously assigned to *K. nataicola*.

The NCIB 8034 strain, historically identified as *Gluconacetobacter xylinus*, matches most closely with *K. oboediens* by phylogenetic analysis in the protein space. This result is confirmed by an ANI of 98.6% to the *K. oboediens* IMBG 311 strain (Error! Reference source not found.). The next nearest neighbor by ANI is *K. intermedius* AF2 at 94.1%, and the ANI to the representative strain of the historical taxon, *K. xylinus* DSM 2325, is 84.6%. The NCIB 8034 strain can unambiguously be reassigned from *G. xylinus* to *K. oboediens*.

The NQ5 strain, historically identified as *G. hansenii*, matches most closely with *N. hansenii* by phylogenetic analysis in the protein space. This result is confirmed by an ANI of 98.9% to the *N. hansenii* C110 strain (Error! Reference source not found.). The next nearest neighbor by ANI is *G. entanii* AV429 at 86.6%. The NQ5 strain is therefore confirmed to belong to the same species as the C110 strain, whose genus has been reassigned to *Novacetimonas*. The NQ5 strain should therefore be taxonomically identified as *N. hansenii*. The placement of all of these strains was further verified using the automated computational pipeline “Type (Strain) Genome Server” (TYGS)^54,55^. The whole-genome sequence placements are in agreement with those above, although the name *K. hansenii* is used *in lieu* of *N. hansenii*.

### Genome annotation

The genomes of *K. nataicola* DS12, *K. oboediens* NCIB 8034, and *N. hansenii* NQ5 were then annotated using the Prokaryotic Genome Annotation Pipeline (PGAP) from NCBI^56–58^. If all prokaryotic genomes are annotated with PGAP, the results can be directly compared to one another. Thus, the nearest neighbor genomes *K. nataicola* RZS01, *K. oboediens* IMBG 311, and *N. hansenii* C110 were also annotated with PGAP. Annotation summary metrics are included in **Table 1**. The total number of genes for each species appears comparable to the reference, except for *K. oboediens*, where the number of genes in NCIB 8034 is much higher than in IMGB 311. This is likely due to the fragmented genome of the IMBG 311 assembly. Notably, two CRISPR arrays were annotated in *N. hansenii* NQ5, but no CRISPR arrays were detected in any of the other strains.

**Table 1:** NCBI PGAP Annotation Metrics.

|  | *K. nataicola*<br>DS12 | *K. nataicola*<br>RZS01 | *K. oboediens*<br>NCIB 8034 | *K. oboediens*<br>IMBG 311 | *N. hansenii*<br>NQ5 | *N. hansenii*<br>C110 |
| --- | --- | --- | --- | --- | --- | --- |
| Genes (total) | 3,703 | 3,601 | 3,833 | 3,437 | 3,042 | 3,133 |
| CDSs (total) | 3,627 | 3,523 | 3,757 | 3,386 | 2,968 | 3,056 |
| Genes (coding) | 3,497 | 3,405 | 3,616 | 3,329 | 2,921 | 2,995 |
| CDSs (with protein) | 3,497 | 3,405 | 3,616 | 3,329 | 2,921 | 2,995 |
| Genes (RNA) | 76 | 78 | 76 | 51 | 74 | 77 |
| rRNAs | 5, 5, 5<br>(5S, 16S, 23S) | 5, 5, 5<br>(5S, 16S, 23S) | 5, 5, 5 (5S, 16S, 23S) | 1, 1, 1 (5S, 16S, 23S) | 5, 5, 5<br>(5S, 16S, 23S) | 5, 5, 5<br>(5S, 16S, 23S) |
| complete rRNAs | 5, 5, 5<br>(5S, 16S, 23S) | 5, 5, 5<br>(5S, 16S, 23S) | 5, 5, 5 (5S, 16S, 23S) | 1, 1, 1 (5S, 16S, 23S) | 5, 5, 5<br>(5S, 16S, 23S) | 5, 5, 5<br>(5S, 16S, 23S) |
| tRNAs | 57 | 59 | 57 | 44 | 55 | 58 |
| ncRNAs | 4 | 4 | 4 | 4 | 4 | 4 |
| Pseudo Genes (total) | 130 | 118 | 141 | 57 | 47 | 61 |
| CDSs (without protein) | 130 | 118 | 141 | 57 | 47 | 61 |
| Pseudo Genes (ambiguous residues) | 0 of 130 | 0 of 118 | 0 of 141 | 0 of 57 | 0 of 47 | 0 of 61 |
| Pseudo Genes (frameshifted) | 66 of 130 | 57 of 118 | 56 of 141 | 29 of 57 | 20 of 47 | 19 of 61 |
| Pseudo Genes (incomplete) | 90 of 130 | 81 of 118 | 95 of 141 | 34 of 57 | 29 of 47 | 48 of 61 |
| Pseudo Genes (internal stop) | 24 of 130 | 21 of 118 | 23 of 141 | 14 of 57 | 11 of 47 | 12 of 61 |
| Pseudo Genes (multiple problems) | 42 of 130 | 36 of 118 | 30 of 141 | 17 of 57 | 10 of 47 | 15 of 61 |
| CRISPR Arrays |  |  |  |  | 2 |  |

### Organization of bcs operons

The production of bacterial cellulose in the *Komagataeibacter* and *Novacetimonas* genera is controlled by bacterial cellulose synthase (*bcs*) genes in several operons. The organization of type I^6^ and type II^22^ operons have been clearly defined. The organization of operons in the present genomes was determined and is depicted in **Figure 6**.

**Figure 6:**
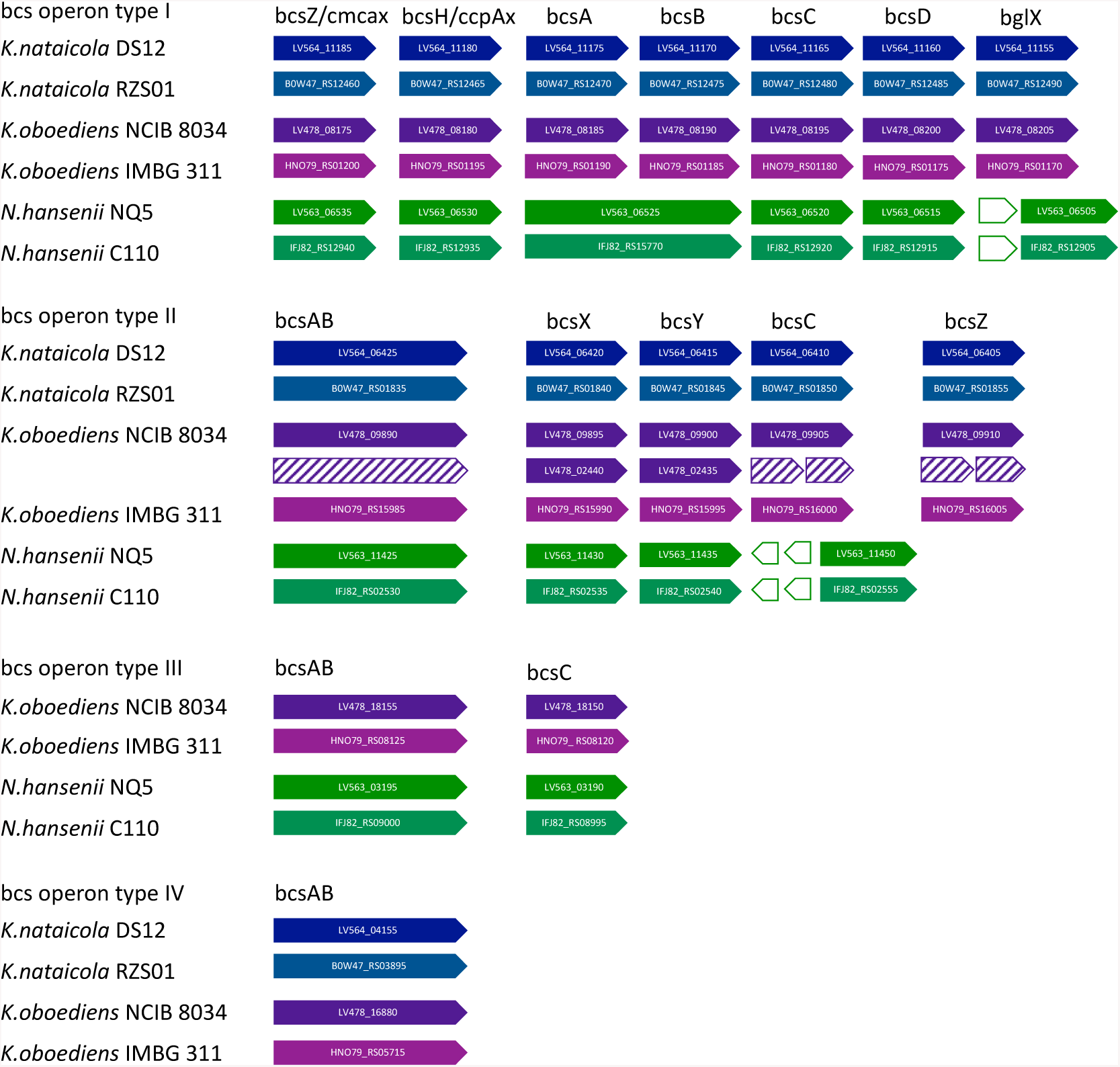
Four types of bcs operons have been reported within the literature. All species studied here had type I and II operons, with K. oboediens NCIB 8034 containing a partial second allele. K. oboediens NCIB 8034 and N. hansenii NQ5 both have type III operons, and K.nataicola DS12 and K.oboediens NCIB 8034 both contained copies of bcsAB not associated with other bcs proteins, here referred to as a type IV operon.

We sought to analyze the *bcs* operon organization in comparison to the type strains for each species and the reference genomes. First, the structure of the operons in *K. nataicola* DS12 were analyzed. Although the genome for the type strain of *K. nataicola* (LMG 1536^59^) is not currently available, we again used the NCBI representative genome of *K. nataicola* (RZS01), which is complete^60^. The genome of RZS01 is reported to contain two *bcs* operons of types I and II. We report the same *bcs* type I operon containing *bcsZ* (also called *cmcAX*), *bcsH* (also called *ccpAX*), *bcsA*, *bcsB*, *bcsC*, *bcsD*, and *bgIxA* (LV564_11185-LV564_11155). The DS12 *bcs* type 2 operon contains *bcsAB*, *bcsX*, *bcsY*, *bcsC*, and *bcsZ* (LV564_06930-LV564_06910). The *bcsZ* gene is not reported as part of the bcs type 2 operon in RZS01, but a homologous gene (B0W47_RS01855) was identified immediately downstream of the bcsC gene (B0W47_RS01850). The *bcsC* gene in the type II operon is entire, and is not disrupted by an insertion sequence as in the *K. xylinus* E25 genome^22^. A third *bcsAB* gene is present in both DS12 (LV564_04155) and RSZ01 (B0W47_RS03895). This gene was not associated with a *bcsC* gene, and is therefore designated as type IV based upon previously-described operon grouping hierarchies^7–9^. The organization of the *bcs* operons in *K. nataicola* DS12 and RSZ01 are therefore identical. The annotators of the RSZ01 genome report the absence of a *pilZ* catalytic domain in the operon II and IV *bcsAB* genes (*bcsABII*, B0W47_RS01835 and *bcsABIII*, B0W47_RS03895)^60^. We report the presence of a *pilZ* catalytic domain in amino acid 574-679 of *bcsAI*, 555-653 of *bcsABII* and 555-653 of *bcsABIII* for both genomes by InterPro sequence search (IPR009875). Based on this analysis, we expect all copies of *bcsA* to be catalytically active in *K. nataicola*.

Next, the bcs operons in *N. hansenii* NQ5 were analyzed, and found to be arranged as previously reported^6,9,22,61^. The type I operon contained *bcsZ*, *bcsH*, *bcsAB*, *bcsC*, *bcsD*, an unknown hypothetical protein, and *bglX* (LV563_06535-06505). Species of the Novacetimonas genera have combined *bcsAB* genes within the type I operon^9^. The second operon contained *bcsAB*, *bcsX*, *bcsY*, two hypothetical proteins (in complement) and *bcsCII* (LV563_11425-11450). No *bcsZ* gene was found in the second operon, as is typical for Novacetimonas. The third operon contained *bcsAB* and *bcsC* (LV563_03195 and 03190).

Finally, the *K. oboediens* NCIB 8034 genome was found to contain five bcs operons, representing all four operon types reported in the reference strain IMBG 311^62^. The first operon contains *bcsZ*, *bcsH*, *bcsA*, *bcsB*, *bcsC*, *bcsD*, and *bglX* (LV478_08175-08205). Two copies of the second operon type are present, *bcs2A* containing *bcsAB*, *bcsX*, *bcsY*, *bcsCII*, and *bcsZ* (LV478_09890-09910) and *bcs2B* containing a frameshifted *bcsAB* pseudogene, *bcsX*, *bcsY*, a disrupted copy of *bcsC*, and a disrupted copy of *bcsZ* (LV478_02445-02415). Multiple copies of the bcs2 operon in *K. oboediens* have been previously reported, with the 174Bp2 strain containing four copies^6^. Disruption of the *bcsC* type II operon gene is also reported in the *K. xylinus* E25 strain as well as other related species^22^. The third operon type comprised *bcsAB* and *bcsC* (LV478_18155 and 18150). The fourth operon type was represented by *bcsAB* alone (LV478_16880). It is interesting to note that the number of operons did not correlate with the highest BNC productivity.

### Presence of CRISPR arrays in *N. hansenii*

As previously described^6^, CRISPR arrays of the type 1-E were found in the genome of *N. hansenii* NQ5. The CRISPR elements *cas3*, *casABCDE*, *cas1* and *cas2* were found adjacent to one another (LV563_ LV563_00855-895). Notably, the *cas3* gene was split into two ORFs, with the upstream ORF (LV563_00855) containing the *cas3* HD endonuclease domain (InterPro domain IPR006483) and the downstream ORF (LV563_860) containing the *cas3* helicase domain (InterPro IPR006474). It is unknown whether the *cas3* gene is therefore active in this strain. A potential explanation for this disruption is the presence of an IS5 family transposase immediately upstream of the *cas3* gene (LV563_850). Orthologous gene clusters were found in *K. medellinensis*, *K. melomenusus*, *N. maltaceti*, and *N. pomaceti*, but not the C110 reference strain of *N. hansenii*. No homologs to the NQ5 *cas3* gene were found in the C110 genome. CRISPR arrays were also not found in *K. nataicola* DS12 or *K. oboediens* NCIB 8034. No known CRISPR arrays have been reported in *K. nataicola* strains, however *K. oboediens* IMBG 180 is reported to contain two CRISPR arrays. These results indicate that CRISPR arrays may not be a highly conserved feature in either *Komagataeibacter* or *Novacetimonas* genomes.

### Conservation of insertional sequences IS1031 and IS1182

Based on a comprehensive analysis of mobile genetic elements^6^, a targeted search for the most notorious insertional sequences within the bacterial group was carried out. Irreversible inactivation of the cellulose producing operons, resulting in cultures dominated by “cel-“ mutants, has long been attributed to the IS5 family insertional sequence IS1031, first identified in *N. hansenii* NCIB 8246 (ATCC 23769)^63,64^. The protein sequence for this transposon (accession AAA25029.1) was blasted against the novel genomes, yielding several hits with high identity and coverage. All hits were found to be contained within orthogroup OG0000002 from the OrthoFinder analysis, and nearly every species included in the analysis contained an ortholog within this orthogroup (Error! Reference source not found.). Many species contained several orthologs, with *K. xylinus* DSM 2325 containing the most, at 12. Out of the genomes generated in this study, *K. oboediens* NCIB 8034 contained the most orthologs at nine, perhaps explaining the previously noted operon duplications and inactivations. *K. nataicola* DS12 contained six orthologs, and *N. hansenii* NQ5 contained only a single ortholog.

Interestingly, though IS1031 orthologs were highly overrepresented in *K. xylinus* DSM 2325, a recent study identified that the insertional sequence most responsible for the spontaneous generation of cel-mutants was an insertional sequence of the IS1182 family^65^. A mutation in the bcsA gene was introduced, which provided resistance to IS1182 insertion and boosted cellulose production. This family of insertion sequences was correlated with orthogroup OG0000170 (Error! Reference source not found.). This orthogroup was well-represented among the species included in the analysis, although fewer orthologs per species were included that for the IS1031 orthogroup. The *K. nataicola* DS12 strain had one ortholog identified, and neither *K. oboediens* NCIB 8034 nor *N. hansenii* NQ5 had a single ortholog placed.

## DISCUSSION

The genomes presented here are complete, contain manually curated annotations for the *bcs* and *cas* operons, and are accompanied by phenotypic data. The metrics presented here indicate that the genomes are of comparable quality to NCBI representative genomes for the species and are of acceptable quality for genomics studies. Furthermore, the *N. hansenii* NQ5 assembly offers a significant improvement in structure over the existing *N. hansenii* genomes, which are highly fragmented into many small contigs.

The architecture of the *bcs* operons for *K. nataicola* DS12 were largely in agreement with the operons reported for the RZS01 strain, providing evidence for conservation of cellulose synthesis machinery across the species. The operon arrangement for the *N. hansenii* NQ5 strain has been well-documented in the past, and our analysis supports previous reports. In contrast, the *bcs* operon arrangement in *K. oboediens* appears to be less stable, with operon duplications common and observed heterogeneity between strains. This note is particularly salient as the NCIB 8034 strain, which was previously identified as a strain of *K. xylinus* and has been used in metabolic engineering studies for cellulose production, is actually a strain of *K. oboediens*. The genomic evidence presented here should be useful to further engineering efforts using this strain or other strains of this species, especially if genomic manipulation of the *bcs* operons is to be undertaken.

Bacterial acquired immunity to phage and other pathogens via CRISPR-Cas systems has been explored in great detail over the last decade^66^. While CRISPR systems are common in bacteria, they are not universal. Indeed, among the three genomes presented here, only *N. hansenii* NQ5 was annotated with a CRISPR array, which may be nonfunctional due to a disrupted *cas3* gene. The absence of CRISPR-cas systems in *K. nataicola* DS12 and K*. oboediens* NCIB 8034, as well as the absence of *cas3* orthologs detected within the majority of representative genomes of the *Komagataeibacter* and *Novacetimonas* genera indicates that CRISPR-cas systems are neither common nor well-conserved among bacteria of these genera.

The NQ5 strain of *N. hansenii* is remarkably resistant to the generation of cel-mutants, with researchers working with the strain reporting no cel-mutants arising over the course of 20 years working with the strain^10^. This makes the strain exceedingly amenable to production of cellulose in agitated cultures, a feat which has proven difficult in other strains. The relatively low number of insertional sequences in the strain may explain the genetic stability relative to other cellulose-producing strains. Indeed, the presence of few insertional sequences may be a generalized feature of the species, as few IS were identified in the *N. hansenii* C110 genome in this study as well as in *N. hansenii* NCIB 8246 (ATCC 23769)^6^. Furthermore, it is possible the unique CRISPR arrays of NQ5 may also play a role in phage resistance and stability.

Thus, based on productivity, yield, carbon source flexibility, ability to create transparent cellulose, and apparent genome stability, it is clear that *N. hansenii* NQ5 is an appropriate strain to use in production of optically clear BNC. Furthermore, the approach presented here could be useful in current efforts in the selection of the right host for the right product^67,68^. In that sense, coupled phenotyping and genotyping experiments like those presented here are effective at identifying candidate strains as hosts for bioproduction.

## METHODS

### Strain isolation and culture

A home-brewed kombucha pellicle, originally sourced from Urban Farm (Portland, ME), was streaked onto a black tea agar plate (2 mg/mL steeped black tea was sterile filtered and supplemented with 40 mg/mL sucrose, 10% (v/v) kombucha starter, 15 mg/mL agar, and liquid autoclaved) and allowed to grow at 30°C for 4 days (**Figure 1**). Cellulose production was observed in specific colonies grown on the black tea agar plate. A single cellulose producing colony was picked and re-streaked on a Hestrin-Schramm^69^ media (HS) agar plate (20mg/mL glucose, 5 mg/mL peptone, 5 mg/mL yeast extract, 1.15 mg/mL citric acid, 2.7 mg/mL disodium phosphate, 15 mg/mL agar) and allowed to grow at 30°C for an additional 4 days. Single colonies were picked and inoculated into 5 mL of HS medium (20 mg/mL glucose, 5 mg/mL peptone, 5 mg/mL yeast extract, 1.15 mg/mL citric acid, 2.7 mg/mL disodium phosphate) media and allowed to grow on a rotating drum for 4 days at 30°C. Cellulose production ability was confirmed, and the isolated strain was labeled DS-12.

*Komagataeibacter oboediens* NCIB 8034, and *Novacetimonas hansenii* NQ5 cellulose producing strains were purchased from the American Type Culture Collection (ATCC, strains #10245 and 53582, respectively).

### Bacterial growth kinetics

Growth kinetic studies were performed for the three bacterial strains. In 96-well plates, inoculated HS media was supplemented with 0.4% (v/v) cellulase from *Trichoderma reesei* (Sigma-Aldrich). The media was supplemented to 1 mM of each carbon source under investigation [glucose, sucrose, fructose, galactose, mannose, mannitol, xylose, arabinose, arabitol, GlcNAc, and GlcN]. Absorbance measurements were taken at 30 min intervals for 7 d at 600 nm under shaken conditions at 30°C (Biotek Synergy H1, Winooski, VT). The maximum growth rate was identified from the linear region of the kinetic curves.

### Pellicle production and cellulose yield from alternative carbon sources

Each bacterial strain was streaked onto HS agar plates, and an individual colony picked. The picked colony was inoculated into 5 mL of Hestrin Schramm (HS) medium and incubated for 4 days on a shaker at 30°C. HS medium was supplemented with 1 mM of glucose for *K. nataicola* and *N. hansenii*, or mannitol for *K. oboediens*. Agitated cultures were then expanded in static cultures using 10% (v/v) inoculum in fresh HS medium.

Previously described methods for cellulose production and yield were used^44^. In brief, fresh HS media was inoculated with 0.1% (v/v) static cultured bacteria and supplemented to 1 mM of each carbon source under investigation [glucose, sucrose, fructose, galactose, mannose, mannitol, xylose, arabinose, arabitol, GlcNAc, and GlcN]. To allow for pellicle formation each bacteria carbon source mixture was cultured in 12-well plates under static conditions at 30°C for 7 d. Pellicles were then harvested, washed in 0.1 M NaOH, rinsed with DI H_2_O, and then lyophilized using a shelf lyophilizer (Labconco, Kansas City, MO). Lyophilized pellicles were weighed, and cellulose yield was calculated relative to weight of carbon source initially supplemented into the culture using the following equation:

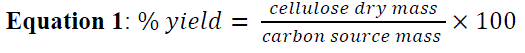

### Cellulose Light Transmission Spectroscopy Measurements

BC pellicles were measured for overall light transmission in the visible light region using a UV-Vis spectrophotometer (Thermo Scientific Evolution 300 UV-Visible Spectrophotometer, Waltham, MA) for pellicles produced by all three bacterial strains. Transmission spectral readings were performed between wavelengths of 300-800 nm with a step size of 2 nm and the visible light transmission (400 – 700 nm range) is reported here.

### Genomic DNA Isolation

Single colonies of the DS-12 isolate, *K. oboediens* NCIB 8034, and *N. hansenii* NQ5 were picked into 5mL HS media with 0.2% (v/v) cellulase from *Trichoderma reesei* (Sigma-Aldrich C2730) and grown to stationary phase. Genomic DNA was isolated from the DS-12 isolate using the Wizard gDNA Purification Kit (Promega A1125) per the manufacturer’s instructions. Genomic DNA from *K. oboediens* NCIB 8034 and *N. hansenii* NQ5 was isolated using the Monarch HMW DNA Extraction Kit for Tissue (NEB T3060S) per the manufacturer’s instructions. The resulting genomic DNA concentration was quantified using the Qubit dsDNA broad range assay (ThermoFisher Q32850).

### Whole Genome Sequencing

Genomic DNA samples were prepared for long-read sequencing using the Rapid Sequencing Kit (Oxford Nanopore SQK-RAD004). The resulting libraries were sequenced using a MinION flow cell (ONT FLO-MIN106D). Genomic DNA samples were prepared for short-read sequencing using the Illumina DNA Prep kit (Illumina 20018704). Resulting libraries were sequenced using an Illumina iSeq 100 sequencer (2×150 paired end reads).

### Read processing

Oxford Nanopore basecalling was performed in real time using the default Guppy basecalling package called by MinKnow. Barcode demultiplexing and trimming was also performed using default MinKnow settings. Illumina reads were basecalled, demultiplexed and trimmed using Illumina iSeq 100 Real-Time Analysis software with default settings. Illumina reads were analyzed with FastQC v0.11.9. All read sets passed “per base sequence quality” metrics.

Nanopore read coverage for the NQ5 strain was exceptionally high, resulting in computational stalling during the genome assembly steps. To mitigate this issue, reads were subsampled to approximately 100× coverage using Filtlong (version 0.2.0) with minimum length of 1000 bp and target bases of 3.52×10^8 bp (https://github.com/rrwick/Filtlong). Default parameters were used otherwise. All raw Nanopore reads were used for assembly of the DS12 and NCIB 8034 genomes.

### Genome assembly and annotation

Genome assembly was carried out on the WPI High-Performance Computing Turing server. Processed Nanopore reads were assembly using Flye^70^ version 2.9-b1768, flags – threads 16, other parameters default. Resultant assemblies were polished with Nanopore reads using Medaka (Oxford Nanopore Technologies) version 1.4.4, flags –t 16 –m r941_min_fast_g303, other parameters default. Illumina reads were aligned to this assembly using Bowtie2^71,72^ version 2.2.5 in paired-end mode with default options. The output.sam file was converted to a.bam file, sorted and indexed using SAMtools^73^ version 1.13. Polished assemblies were further polished with Illumina paired-end reads, using Pilon^74^ version 1.24, flags –Xmx64G, other flags default.

Putative plasmids were identified by the presence of a repA homolog from closely related species, using methodology and accession numbers described by Ryngajłło et al^6^. Contigs containing neither a repA gene nor a dnaA gene were scrutinized for the presence of meaningful annotations, and discarded as assembly artifacts if no meaningful annotations were found. Chromosomal contigs were identified both by the presence of a dnaA gene and by size (largest contig). Contigs were renamed as numbered plasmids or chromosomes.

Genome assembly summary metrics were calculated using QUAST version 5.2.0 with flags –-space-efficient –-no-plots –-no-html –-no-icarus. Genome completeness was assessed using BUSCO version 5.0.0, with flags –l rhodospirillales_odb10 –m genome, and other settings default. Assemblies were benchmarked against representative genomes accessed from the NCBI Genome database on 1/8/2023. The representative genomes were as follows: *K. nataicola*, strain RZS01, accession GCF_002009295.1; *K. oboediens*, strain IMBG 311, GCF_018491655.1; *N. hansenii*, strain C110, accession GCF_014843995.1.

Genome assemblies were submitted to the National Center for Biotechnology Information (NCBI) and annotated with the NCBI Prokaryotic Genome Annotation Pipeline (PGAP)^56–58^ under the BioProject ID PRJNA789846.

### Phylogenetic placement

Protein sequences from representative genomes of all species in the *Gluconacetobacter*, *Komagataeibacter* and *Novacetimonas* genera were accessed from RefSeq (Error! Reference source not found.). The OrthoFinder^75,76^ tool was used to construct a phylogenetic tree of these species based upon OrthoGroups shared between the species. The IcyTree^77^ tool was used to draw the graph. Nucleotide sequences for the genomes of species in the Komagataeibacter and Novacetimonas genera in Error! Reference source not found. were downloaded from RefSeq. The ANI Genome-based distance matrix calculator^52^ was used to calculate the ANI between all genomes provided.

### bcs operon analysis

Genomic annotations for each strain were compared to annotations for the *Komagataeibacter xylinus* E25 strain^6,22^. Each variant of the bcsA gene was aligned to the novel genomes using NCBI BLAST. Copies of the bcsA genes were identified and the genomic loci were noted. Additional genes in each operon were identified by comparison to the genes in the *K. xylinus* E25 genome and by analyzing consecutive features within the annotated genome.

## STATEMENT OF AUTHOR ROLES

EMvZ, KWK, JMC, and EMY conceived of the study. JMC supervised the phenotypic characterization. EMY supervised the genomics. EMvZ isolated DS12 and identified NQ5 and NCIB 8034 as strains of interest and performed phenotypic characterization experiments. JHC extracted genomic DNA and performed ONT and Illumina sequencing of DS12. KWK extracted genomic DNA and performed ONT and Illumina sequencing of NQ5 and NCIB 8034 strains, assembled genomes and performed comparative genomics. KWK and EMvZ contributed text and figures to the paper. CN assisted with phylogenetic analysis. SJW provided genomics guidance.

## ABBREVIATIONS

ANI: average nucleotide identity; BNC: bacterial nanocellulose; NCBI: National Center for Biotechnology Information; PGAP: prokaryotic genome annotation pipeline

## DATA AVAILABILITY

This Whole Genome Shotgun project has been deposited at DDBJ/ENA/GenBank under the accession PRJNA789846. Raw ONT and Illumina sequencing reads (SRA submission) can be accessed under this BioProject accession. Complete genomes were submitted to GenBank and can be accessed as genomes associated with this BioProject, or by individual accessions: *K. nataicola* DS12 CP118672-CP118676, *K. oboediens* NCIB 8034 CP117858-CP117864, *N. hansenii* NQ5 CP117863-CP117864.

## REFERENCES

1. Chandana, A., Mallick, S. P., Dikshit, P. K., Singh, B. N. & Sahi, A. K. Recent Developments in Bacterial Nanocellulose Production and its Biomedical Applications. J. Polym. Environ. 30, 4040– 4067 (2022).

2. Huang, Y. et al. Flexible cathodes and multifunctional interlayers based on carbonized bacterial cellulose for high-performance lithium–sulfur batteries. J. Mater. Chem. A 3, 10910–10918 (2015).

3. Ludwicka, K., Kaczmarek, M. & Białkowska, A. Bacterial Nanocellulose—A Biobased Polymer for Active and Intelligent Food Packaging Applications: Recent Advances and Developments. Polymers 12, 2209 (2020).

4. Bae, S. & Shoda, M. Bacterial Cellulose Production by Fed-Batch Fermentation in Molasses Medium. Biotechnol. Prog. 20, 1366–1371 (2004).

5. Keshk, S. & Sameshima, K. The utilization of sugar cane molasses with/without the presence of lignosulfonate for the production of bacterial cellulose. Appl. Microbiol. Biotechnol. 72, 291–296 (2006).

6. Ryngajłło, M., Kubiak, K., Jędrzejczak-Krzepkowska, M., Jacek, P. & Bielecki, S. Comparative genomics of the Komagataeibacter strains—Efficient bionanocellulose producers. MicrobiologyOpen 8, e00731 (2019).

7. Gullo, M., La China, S., Petroni, G., Di Gregorio, S. & Giudici, P. Exploring K2G30 Genome: A High Bacterial Cellulose Producing Strain in Glucose and Mannitol Based Media. Front. Microbiol. 10, (2019).

8. Liu, M. et al. Complete genome analysis of Gluconacetobacter xylinus CGMCC 2955 for elucidating bacterial cellulose biosynthesis and metabolic regulation. Sci. Rep. 8, 6266 (2018).

9. Brandão, P. R., Crespo, M. T. B. & Nascimento, F. X. Phylogenomic and comparative analyses support the reclassification of several Komagataeibacter species as novel members of the Novacetimonas gen. nov. and bring new insights into the evolution of cellulose synthase genes. Int. J. Syst. Evol. Microbiol. 72, 005252 (2022).

10. Czaja, W., Romanovicz, D. & Brown, R. malcolm. Structural investigations of microbial cellulose produced in stationary and agitated culture. Cellulose 11, 403–411 (2004).

11. Liu, L. et al. Komagataeibacter cocois sp. nov., a novel cellulose-producing strain isolated from coconut milk. Int. J. Syst. Evol. Microbiol. 68, 3125–3131 (2018).

12. Slapšak, N., Cleenwerck, I., De Vos, P. & Trček, J. Gluconacetobacter maltaceti sp. nov., a novel vinegar producing acetic acid bacterium. Syst. Appl. Microbiol. 36, 17–21 (2013).

13. Škraban, J., Cleenwerck, I., Vandamme, P., Fanedl, L. & Trček, J. Genome sequences and description of novel exopolysaccharides producing species Komagataeibacter pomaceti sp. nov. and reclassification of Komagataeibacter kombuchae (Dutta and Gachhui 2007) Yamada et al., 2013 as a later heterotypic synonym of Komagataeibacter hansenii (Gosselé et al. 1983) Yamada et al., 2013. Syst. Appl. Microbiol. 41, 581–592 (2018).

14. Marič, L., Cleenwerck, I., Accetto, T., Vandamme, P. & Trček, J. Description of Komagataeibacter melaceti sp. nov. and Komagataeibacter melomenusus sp. nov. Isolated from Apple Cider Vinegar. Microorganisms 8, E1178 (2020).

15. Iino, T. et al. Gluconacetobacter kakiaceti sp. nov., an acetic acid bacterium isolated from a traditional Japanese fruit vinegar. Int. J. Syst. Evol. Microbiol. 62, 1465–1469 (2012).

16. Castro, C. et al. Gluconacetobacter medellinensis sp. nov., cellulose– and non-cellulose-producing acetic acid bacteria isolated from vinegar. Int. J. Syst. Evol. Microbiol. 63, 1119–1125 (2013).

17. Naloka, K., Yukphan, P., Matsutani, M., Matsushita, K. & Theeragool, G. Komagataeibacter diospyri sp. nov., a novel species of thermotolerant bacterial nanocellulose-producing bacterium. Int. J. Syst. Evol. Microbiol. 70, 251–258 (2020).

18. R, R., et al. Bacterial nanocellulose: engineering, production, and applications. Bioengineered 12, 11463–11483.

19. de Amorim, J. D. P. et al. Plant and bacterial nanocellulose: production, properties and applications in medicine, food, cosmetics, electronics and engineering. A review. Environ. Chem. Lett. 18, 851–869 (2020).

20. Florea, M. et al. Engineering control of bacterial cellulose production using a genetic toolkit and a new cellulose-producing strain. Proc. Natl. Acad. Sci. 113, E3431–E3440 (2016).

21. Omadjela, O. et al. BcsA and BcsB form the catalytically active core of bacterial cellulose synthase sufficient for in vitro cellulose synthesis. Proc. Natl. Acad. Sci. U. S. A. 110, 17856–17861 (2013).

22. Szymczak, I., Pietrzyk-Brzezińska, A. J., Duszyński, K. & Ryngajłło, M. Characterization of the Putative Acylated Cellulose Synthase Operon in Komagataeibacter xylinus E25. Int. J. Mol. Sci. 23, 7851 (2022).

23. Yadav, V. et al. Novel In Vivo-Degradable Cellulose-Chitin Copolymer from Metabolically Engineered Gluconacetobacter xylinus. Appl. Environ. Microbiol. 76, 6257–6265 (2010).

24. Collins, J. H. et al. Engineered yeast genomes accurately assembled from pure and mixed samples. Nat. Commun. 12, 1485 (2021).

25. Keating, K. & Young, E. M. Systematic part transfer by extending a modular toolkit to diverse bacteria. 2023.02.07.527528 Preprint at 10.1101/2023.02.07.527528 (2023).

26. Jang, W. D. et al. Genomic and metabolic analysis of Komagataeibacter xylinus DSM 2325 producing bacterial cellulose nanofiber. Biotechnol. Bioeng. 116, 3372–3381 (2019).

27. Bureau, T. E. & Brown, R. M. In vitro synthesis of cellulose II from a cytoplasmic membrane fraction of Acetobacter xylinum. Proc. Natl. Acad. Sci. 84, 6985–6989 (1987).

28. Kazuwo, M., et al. Method for producing 2-keto-lgulonic acid. (1966).

29. Schramm, M., Gromet, Z. & Hestrin, S. Synthesis of cellulose by Acetobacter xylinum. 3. Substrates and inhibitors. Biochem. J. 67, 669–679 (1957).

30. Wang, S.-S., et al. Insights into Bacterial Cellulose Biosynthesis from Different Carbon Sources and the Associated Biochemical Transformation Pathways in Komagataeibacter sp. W1. Polymers 10, 963 (2018).

31. Molina-Ramírez, C., et al. Effect of Different Carbon Sources on Bacterial Nanocellulose Production and Structure Using the Low pH Resistant Strain Komagataeibacter Medellinensis. Materials 10, 639 (2017).

32. Tyagi, N. & Suresh, S. Production of cellulose from sugarcane molasses using Gluconacetobacter intermedius SNT-1: optimization & characterization. J. Clean. Prod. 112, 71–80 (2016).

33. Lin, S.-P., et al. Isolation and identification of cellulose-producing strain Komagataeibacter intermedius from fermented fruit juice. Carbohydr. Polym. 151, 827–833 (2016).

34. Mohammadkazemi, F., Azin, M. & Ashori, A. Production of bacterial cellulose using different carbon sources and culture media. Carbohydr. Polym. 117, 518–523 (2015).

35. Trovatti, E., Serafim, L. S., Freire, C. S. R., Silvestre, A. J. D. & Neto, C. P. Gluconacetobacter sacchari: An efficient bacterial cellulose cell-factory. Carbohydr. Polym. 86, 1417–1420 (2011).

36. Son, H.-J., Heo, M.-S., Kim, Y.-G. & Lee, S.-J. Optimization of fermentation conditions for the production of bacterial cellulose by a newly isolated Acetobacter. Biotechnol. Appl. Biochem. 33, 1–5 (2001).

37. Fang, L. & Catchmark, J. M. Characterization of cellulose and other exopolysaccharides produced from Gluconacetobacter strains. Carbohydr. Polym. 115, 663–669 (2015).

38. Lee, J. W. et al. Direct Incorporation of Glucosamine andN-Acetylglucosamine into Exopolymers by Gluconacetobacter xylinus (=Acetobacter xylinum) ATCC 10245: Production of Chitosan-Cellulose and Chitin-Cellulose Exopolymers. Appl. Environ. Microbiol. 67, 3970–3975 (2001).

39. Malik, S., Singh, M. & Mathur, A. Antimicrobial Activity of Food Grade Glucosamine.

40. Antolak, H., Piechota, D. & Kucharska, A. Kombucha Tea—A Double Power of Bioactive Compounds from Tea and Symbiotic Culture of Bacteria and Yeasts (SCOBY). Antioxidants 10, 1541 (2021).

41. Zhang, H. et al. Reconstruction of a Genome-scale Metabolic Network of Komagataeibacter nataicola RZS01 for Cellulose Production. Sci. Rep. 7, 7911 (2017).

42. van Zyl, E. M. & Coburn, J. M. Hierarchical structure of bacterial-derived cellulose and its impact on biomedical applications. Curr. Opin. Chem. Eng. 24, 122–130 (2019).

43. La China, S. et al. Genome sequencing and phylogenetic analysis of K1G4: a new Komagataeibacter strain producing bacterial cellulose from different carbon sources. Biotechnol. Lett. 42, 807–818 (2020).

44. van Zyl, E. M. et al. Structural properties of optically clear bacterial cellulose produced by Komagataeibacter hansenii using arabitol. Biomater. Adv. 148, 213345 (2023).

45. Jeannine Coburn, Elzani van Zyl, & Eric Young. Cellulose-Based Materials and Methods of Making the Same.

46. Gurevich, A., Saveliev, V., Vyahhi, N. & Tesler, G. QUAST: quality assessment tool for genome assemblies. Bioinformatics 29, 1072–1075 (2013).

47. Meier-Kolthoff, J. P., Klenk, H.-P. & Göker, M. Taxonomic use of DNA G+C content and DNA-DNA hybridization in the genomic age. Int. J. Syst. Evol. Microbiol. 64, 352–356 (2014).

48. Simão, F. A., Waterhouse, R. M., Ioannidis, P., Kriventseva, E. V. & Zdobnov, E. M. BUSCO: assessing genome assembly and annotation completeness with single-copy orthologs. Bioinformatics 31, 3210–3212 (2015).

49. Yamada, Y. et al. Description of *Komagataeibacter* gen. nov., with proposals of new combinations (*Acetobacteraceae*). J. Gen. Appl. Microbiol. 58, 397–404 (2012).

50. Yamada, Y. et al. Subdivision of the genus Gluconacetobacter Yamada, Hoshino and Ishikawa 1998: the proposal of Komagatabacter gen. nov., for strains accommodated to the Gluconacetobacter xylinus group in the α-Proteobacteria. Ann. Microbiol. 62, 849–859 (2012).

51. Jelenko, K., Cepec, E., Nascimento, F. X. & Trček, J. Comparative Genomics and Phenotypic Characterization of Gluconacetobacter entanii, a Highly Acetic Acid-Tolerant Bacterium from Vinegars. Foods 12, 214 (2023).

52. Rodriguez-R, L. M. & Konstantinidis, K. T. The enveomics collection: a toolbox for specialized analyses of microbial genomes and metagenomes. https://peerj.com/preprints/1900 (2016) doi:10.7287/peerj.preprints.1900v1.

53. Ciufo, S. et al. Using average nucleotide identity to improve taxonomic assignments in prokaryotic genomes at the NCBI. Int. J. Syst. Evol. Microbiol. 68, 2386–2392 (2018).

54. Meier-Kolthoff, J. P. & Göker, M. TYGS is an automated high-throughput platform for state-of-the-art genome-based taxonomy. Nat. Commun. 10, 2182 (2019).

55. Meier-Kolthoff, J. P., Carbasse, J. S., Peinado-Olarte, R. L. & Göker, M. TYGS and LPSN: a database tandem for fast and reliable genome-based classification and nomenclature of prokaryotes. Nucleic Acids Res. 50, D801–D807 (2022).

56. Li, W. et al. RefSeq: expanding the Prokaryotic Genome Annotation Pipeline reach with protein family model curation. Nucleic Acids Res. 49, D1020–D1028 (2021).

57. Haft, D. H. et al. RefSeq: an update on prokaryotic genome annotation and curation. Nucleic Acids Res. 46, D851–D860 (2018).

58. Tatusova, T. et al. NCBI prokaryotic genome annotation pipeline. Nucleic Acids Res. 44, 6614–6624 (2016).

59. Lisdiyanti, P., Navarro, R. R., Uchimura, T. & Komagata, K. Reclassification of Gluconacetobacter hansenii strains and proposals of Gluconacetobacter saccharivorans sp. nov. and Gluconacetobacter nataicola sp. nov. Int. J. Syst. Evol. Microbiol. 56, 2101–2111 (2006).

60. Zhang, H. et al. Complete genome sequence of the cellulose-producing strain Komagataeibacter nataicola RZS01. Sci. Rep. 7, 4431 (2017).

61. Florea, M., Reeve, B., Abbott, J., Freemont, P. S. & Ellis, T. Genome sequence and plasmid transformation of the model high-yield bacterial cellulose producer Gluconacetobacter hansenii ATCC 53582. Sci. Rep. 6, 23635 (2016).

62. Orlovska, I. et al. Bacterial Cellulose Retains Robustness but Its Synthesis Declines After Exposure to a Mars-like Environment Simulated Outside the International Space Station. Astrobiology 21, 706–717 (2021).

63. Coucheron, D. H. An Acetobacter xylinum insertion sequence element associated with inactivation of cellulose production. J. Bacteriol. 173, 5723–5731 (1991).

64. Coucheron, D. H. A family of IS1031 elements in the genome of Acetobacter xylinum: nucleotide sequences and strain distribution. Mol. Microbiol. 9, 211–218 (1993).

65. Hur, D. H. et al. Enhanced production of cellulose in Komagataeibacter xylinus by preventing insertion of IS element into cellulose synthesis gene. Biochem. Eng. J. 156, 107527 (2020).

66. Nussenzweig, P. M. & Marraffini, L. A. Molecular Mechanisms of CRISPR-Cas Immunity in Bacteria. Annu. Rev. Genet. 54, 93–120 (2020).

67. Collins, J. H. & Young, E. M. Genetic engineering of host organisms for pharmaceutical synthesis. Curr. Opin. Biotechnol. 53, 191–200 (2018).

68. Keating, K. W. & Young, E. M. Synthetic biology for bio-derived structural materials. Curr. Opin. Chem. Eng. 24, 107–114 (2019).

69. Schramm, M. & Hestrin, S. Synthesis of cellulose by Acetobacter xylinum. 1. Micromethod for the determination of celluloses*. Biochem. J. 56, 163–166 (1954).

70. Kolmogorov, M., Yuan, J., Lin, Y. & Pevzner, P. A. Assembly of long, error-prone reads using repeat graphs. Nat. Biotechnol. 37, 540–546 (2019).

71. Langmead, B. & Salzberg, S. L. Fast gapped-read alignment with Bowtie 2. Nat. Methods 9, 357– 359 (2012).

72. Langmead, B., Wilks, C., Antonescu, V. & Charles, R. Scaling read aligners to hundreds of threads on general-purpose processors. Bioinformatics 35, 421–432 (2019).

73. Danecek, P. et al. Twelve years of SAMtools and BCFtools. GigaScience 10, giab008 (2021).

74. Walker, B. J. et al. Pilon: An Integrated Tool for Comprehensive Microbial Variant Detection and Genome Assembly Improvement. PLOS ONE 9, e112963 (2014).

75. Emms, D. M. & Kelly, S. OrthoFinder: solving fundamental biases in whole genome comparisons dramatically improves orthogroup inference accuracy. Genome Biol. 16, 157 (2015).

76. Emms, D. M. & Kelly, S. OrthoFinder: phylogenetic orthology inference for comparative genomics. Genome Biol. 20, 238 (2019).

77. Vaughan, T. G. IcyTree: rapid browser-based visualization for phylogenetic trees and networks. Bioinformatics 33, 2392–2394 (2017).

